# Simulated attack reveals how lesions affect network properties in post-stroke aphasia

**DOI:** 10.1101/2021.11.01.466833

**Authors:** John D. Medaglia, Brian A. Erickson, Dorian Pustina, Apoorva S. Kelkar, Andrew T. DeMarco, J. Vivian Dickens, Peter E. Turkeltaub

**Author notes:** **Author Contributions**, JDM analyzed the data and wrote the manuscript. ASK preprocessed the data and contributed to manuscript writing. PET provided the data, project oversight, and contributed to project conceptualization.

## Abstract

Aphasia is one of the most prevalent cognitive syndromes caused by stroke. The rarity of premorbid imaging and heterogeneity of lesion size and extent obfuscates the links between the local effects of the lesion, global anatomical network organization, and aphasia symptoms. We applied a simulated attack approach to examine the effects of 39 stroke lesions on network topology by simulating their effects in a control sample of 36 healthy brain networks. We focused on measures of global network organization thought to support overall brain function and resilience in the whole brain and within the left hemisphere. After removing lesion volume from the network topology measures and behavioral scores (the Western Aphasia Battery Aphasia Quotient; WAB-AQ), four behavioral factor scores obtained from a neuropsychological battery, and a factor sum), we compared the behavioral variance accounted for by simulated post-stroke connectomes to that observed in the randomly permuted data. Overall, global measures of network topology in the whole brain and left hemisphere accounted for 10% variance or more of the WAB-AQ and the lexical factor score beyond lesion volume and null permutations. Streamline networks provided more reliable point estimates than FA networks. Edge weights and network efficiency were weighted most highly in predicting the WAB-AQ for FA networks. Overall, our results suggest that global network measures can provide modest statistical value predicting overall aphasia severity, but less value in predicting specific behaviors. Variability in estimates could be induced by premorbid ability, deafferentation and diaschisis, and neuroplasticity following stroke.

## Introduction

Aphasia is one of the primary cognitive symptoms following left hemispheric strokes, affecting 180,000 new individuals a year in the United States [1]. Despite decades of research, the brain basis of aphasia outcomes and recovery remain only partially understood. The majority of stroke research has focused on the relationship between the regional anatomical influences of stroke on cognitive symptoms and outcomes [2, 3, 4, 5]. More recently, investigators have studied the relationships between individual anatomical tracts, the topology of complex brain networks (the *connectome*, [6, 7, 8, 9]), and behavior [10, 11, 12, 13, 14].

Post-stroke, the remaining neuroanatomy maintains cognition and supports recovery. Anatomical network connectivity in the lost and residual (spared) connectome after stroke is related to behavior [15, 16, 17, 12, 18, 19, 20, 14]. In particular, single-connection analyses have demonstrated that regions with links to classical *hub* regions such as the temporoparietal junction are crucial for overall language function assessed with clinical measures [14]. Strokes that directly impact network hubs disproportionately lead to global cognitive deficits post-stroke on tasks that place significant semantic or language-production demands on patients [21]. In addition, cognitive outcomes are associated with the preservation of the brain’s modular configuration – the tendency for brain regions to group into well-connected clusters [22, 23]. Overall, these findings suggest that the role of single regions and their connections in network topology, as well as overall network topology, are related to stroke symptomatology.

A primary difficulty in assessing stroke-induced effects on network topology is that researchers often lack premorbid data within-subjects, leading them to rely on cross-sectional analyses. This results in a reference problem for each stroke. Lesions occur within a single subject, but the consequences of the lesion interact with other factors about the individual, such as their development, demographics, and brain organization. As a complement to observing the consequences of stroke and other types of brain injury, “simulated attack” models are computational approaches that apply virtual damage to the brain and measure their putative consequences [24, 25]. These models can be used to systematically quantify the influences of damage to regions and connections on brain network organization. After simulating damage, hypotheses about network robustness, cognitive resilience, and recovery can be tested in the residual connectomes [26]. Measures characterizing the disconnectivity of circuits and networks [12], the overall efficiency of the network [27, 28, 29], and the balance between local and distributed processing (*small-worldness*, [30]) could relate to behavioral performance. In addition, the deviation in these properties from that expected in a comparison model of healthy subjects might also characterize variation in resilience to cognitive decline.

To examine these possibilities in aphasia severity, we used probabalistic diffusion tractography to create anatomical connectomes in 39 subjects with left-hemispheric strokes. Then, we computed measures that quantify five network properties of anatomical connectivity post-stroke thought to be related to the integrity of observed topology. Using a simulated attack model, we computed the effects of each stroke’s specific pattern of connection losses to quantify its effects on the whole brain and intra-left hemisphere connections in a sample of healthy subjects. Then, we computed models estimating the behavioral variance measured with clinical language measures accounted for by simulated anatomical network measures. This technique allowed us to obtain confidence intervals for the strength of brain-behavior relationships between lesioned network topology and behavior. Above and beyond lesion volume, we hypothesized that total edge weights, network modularity, global shortest path length, higher local clustering, and small-worldness would be related to better language performance. We further hypothesized that the global network measures would be more related to global measures of the severity of language deficits than factor scores representing specific lexical, auditory comprehension, phonology, and cognitive/semantic deficits.

## Methods

### Subjects

Data on 61 stroke patients and 37 healthy controls were collected. Twelve patients were excluded from analysis due to missing or abnormal neuroimaging data; 7 due to lesions outside the left hemisphere; one due to missing or at-floor behavioral data; one due to acuteness of stroke; and one due to non-native English language. We excluded 1 healthy subject due to inflated region-wise streamline estimation (twice the connectome edge density as any other subject). After exclusions the final samples consisted of 39 subjects with stroke (mean age = 59.74, St.D. = 9.17, 16 females) and 36 healthy subjects (mean age = 59.13, St.D. = 13.84, 15 females). All subjects were scanned on a 3T Siemens Magnetom scanner at Georgetown University’s Center for Functional and Molecular Imaging (CFMI). Aside from the stroke events in patients, participants had no history of psychiatric or other neurological condition. Healthy controls had no history of neurological disease or developmental disorder. Subjects with stroke had a neurologist confirmed diagnosis of aphasia, a left-hemispheric stroke, and the absence of a right-hemispheric stroke. The data were collected as part of previous studies conducted in the Cognitive Recovery Lab at Georgetown University.

All procedures were approved in a convened review by Georgetown University’s Institutional Review Board and were carried out in accordance with the guidelines of the Institutional Review Board/Human Subjects Committee, Georgetown University. All participants volunteered and provided informed consent in writing prior to data collection.

### Behavioral Data

The Western Aphasia Battery - Revised [31] was obtained for each individual with stroke. In addition, participants with stroke performed a broader battery of tasks previously described in detail [32]. To reduce the scores from the battery, a principal components factor analysis was performed in SPSS 25 using the individual test scores from the WAB-R and the other battery tasks on the 59 participants with stroke who were able to provide complete behavioral data. Factor analysis was performed on the correlation matrix, factors were extracted based on the standard cutoff of eigenvalue > 1, and Varimax rotation with Kaiser normalization was applied to achieve orthogonal factors. Consistent with a previously reported factor analysis on a subset of these participants, the factor analysis revealed 4 factors cumulatively accounting for 83.7% of variance in the scores that we interpreted to reflect lexical production, auditory comprehension, phonology, and cognitive & semantic aspects of behavior (see Table 1). Factor scores for each participant were calculated using the regression method.

**Table 1:** Factor loadings for the behavioral data.

|  | **Factor I (Lexical**<br>**Production)** | **Factor II (Auditory**<br>**Comprehension)** | **Factor III**<br>**(Phonology)** | **Factor IV**<br>**(Cognitive/Semantic)** |
| --- | --- | --- | --- | --- |
| Philadelphia Naming Test | 0.83 | 0.24 | 0.24 | 0.26 |
| WAB Object Naming | 0.79 | 0.36 | 0.33 | 0.21 |
| Reading Real Words | 0.77 | 0.19 | 0.39 | 0.26 |
| Reading word-to-picture matching | 0.75 | 0.39 | 0.32 | 0.11 |
| WAB Spontaneous Speech Fluency | 0.73 | 0.41 | 0.34 | 0.18 |
| WAB Spontaneous Speech Content | 0.7 | 0.4 | 0.31 | 0.31 |
| WAB Responsive Speech | 0.67 | 0.54 | 0.33 | 0.14 |
| WAB Repetition | 0.57 | 0.52 | 0.52 | 0.12 |
| WAB Yes/No Questions | 0.24 | 0.86 | 0.18 | 0.03 |
| WAB Sequential Commands | 0.32 | 0.68 | 0.29 | 0.32 |
| WAB Word Recognition | 0.42 | 0.69 | 0.23 | 0.36 |
| WAB Sentence Completion | 0.58 | 0.62 | 0.3 | 0.12 |
| Digit Span Forwards | 0.29 | 0.43 | 0.76 | 0.21 |
| Digit Span Backwards | 0.45 | 0.16 | 0.72 | 0.31 |
| Pseudoword Repetition | 0.46 | 0.45 | 0.65 | 0.01 |
| Reading Pseudowords | 0.5 | 0.27 | 0.62 | 0.34 |
| Letter Fluency (Total 4 letters) | 0.58 | 0.12 | 0.61 | 0.16 |
| Backward Spatial Span | 0.18 | -0.05 | 0.32 | 0.83 |
| Pyramids and Palm Trees | 0.41 | 0.13 | -0.06 | 0.8 |
| Forward Spatial Span | -0.12 | 0.27 | 0.41 | 0.76 |
| Auditory Word-to-Picture Matching | 0.51 | 0.41 | -0.05 | 0.65 |

### Neuroimaging

Diffusion images were acquired on a Siemens 3.0T Magnetom Trio for all subjects along with a T1-weighted 1mm resolution MPRAGE anatomical scan at each scanning session as part of a larger imaging protocol. We used a high-angular resolution diffusion imaging (HARDI) acquisition scheme with a maximum b-value of 1,100 (80 dirs, 10 b=0; 10 b=300; 60 b = 1100) and a 2.5mm isotropic voxel size. We used a transversal acquisition of 55 axial slices with the following parameters: repetition time (TR) = 7.5 s; echo time (TE) = 87 ms; field of view (FoV) = 240 × 240, 138 mm, matrix = 96, total acquisition time of 10:00. MPRAGE scans were collected with TR = 1900ms, TE = 2.52ms, 176 sagittal slices with 0.9mm slice thickness, FoV = 240 × 240, matrix = 256, inversion time (TI) = 900ms and flip angle = 9°, total acquisition time of 5:34.

#### Anatomical Image Imputation

We conducted full-brain tractography with techniques that reduce tractography artifacts. To achieve this, tractography must be constrained anatomically to seed or terminate streamlines at the grey matter/white matter border [33, 34, 35]. However, identifying tissue types in stroke cases is problematic because of the abnormal signal intensity at the gray-white matter border. We resolved the issue by imputing estimates of healthy tissue.

We imputed anatomical images in two steps. In step 1, lesioned voxels were identified using lesion tracings provided by a experienced cognitive neurologist (coauthor PET). We then flipped the lesioned brain along the left-right plane and registered the flipped brain into onto the non-flipped brain. Next, we filled the lesioned area with the healthy tissue of the homotopic contralesional hemisphere. This procedure can leave visible marks of the filled area due to the sudden change in signal, which may cause artifacts when identifying tissue types. Thus, we imputed a new brain with highly similar morphological features as the original subject’s brain. The imputation procedure used the morphological structure of the reference image (the filled brain from step 1) and the voxel values of a set of healthy control images to produce a new image. Each voxel value in the new image was determined by combining the values from all the healthy images using ANTs’ joint image fusion procedure, where images more similar around the voxel of interest received more weight (similar to multi-altas label fusion, [36]). We conducted a search of the optimal number of healthy brains and the optimal radius of similarity around each voxel to obtain the best result. We obtained an optimal outcome with 22 healthy brains and a radius of 1 (i.e., a single layer of voxels around each voxel is used to check the similarity between images and assign weights to healthy images). We inspected the resulting imputed image to make sure there were no artifacts; none were found. Importantly, the non-lesioned gyri and sulci in the original image followed the gyri and sulci of the imputed image without any visible deviation.

We performed all the imputation procedures in ANTs (v. 2.2.0). Before any processing, all images were skull-stripped (antsBrainExtraction.sh), corrected for magnetic field inhomogneity (N4BiasFieldCorrection), and denoised with an edge preserving algorithm (PeronaMalik, denoising amount: 0.7, iterations: 10). We added back the lesion mask to the brain mask after skull-stripping to ensure that the lesion area was included in the imputation. Each imputation required one registration of the flipped image and 22 registrations of the healthy brains onto the filled image. We conducted all registrations using the SyN non-linear algorithm [37] with cost function masking to remove the lesion mask from consideration during the registration computations [38].

#### Diffusion Tractography

We used MRtrix3 to process the diffusion data [39]. First, we denoised (dwidenoise -extent 9,9,9), corrected for motion and eddy currents (dwipreproc), and corrected for field inhomogneity (dwibiascorrect). Then we computed response functions for multiple tissues using the tissue information available in the DWI data itself (dwi2response dhollander). We finally computed the fiber orientation distribution (FOD) with a multi-shell multi-tissue algorithm (dwi2fod msmt_csd).

To find the GM/WM tissue, we applied tissue classification on the imputed structural image (5ttgen fsl), and then brought the tissue information into DWI space after registering the original (lesioned) T1w image of the subject onto the mean b=0 image (antsRegistration in order: translation, rigid, SyN) and applying the transformations to each tissue type image. We then computed the GM/WM border for use in the next tractography step (5tt2gmwmi). We performed tractography by seeding 15 million streamlines from the GM/WM border (tckgen algorithm: iFOD2, step: 1mm, minlength: 10mm, maxlength: 300mm, angle: 45 degrees, backtrack: crop at gmwmi). We then filtered the tractogram with the SIFT2 algorithm [40, 41] to decrease tractography artifacts. Inter-subject connection density normalization was then achieved through scalar multiplication of each connectome by the subject’s “proportionality coefficient” derived by SIFT2, denoted by *μ*, which represents the estimated fiber volume per unit length contributed by each streamline [41].

#### Network construction

Anatomical scans (imputed in the case of patients) were segmented using FreeSurfer [42] and parcellated using the connectome mapping toolkit [43]. A parcellation scheme including *N* = 234 regions was registered to a single b=0 volume from each subject’s native-space DSI data. The b=0 to MNI voxel mapping produced via Q-Space Diffeomorphic Reconstruction (QSDR) was used to map region labels from native space to MNI coordinates so that individual subject data could be combined and analyzed in a shared standard space. To extend region labels through the grey-white matter interface, the atlas was dilated by 4mm [44]. Dilation was accomplished by filling non-labeled voxels with the statistical mode of their neighbors’ labels. In the event of a tie, one of the modes was randomly selected. Each streamline was labeled according to its terminal region pair. From these data, we constructed an anatomical connectivity adjacency matrix, **A** whose element *A_ij_* represented the average fractional anisotropy (FA) of the streamlines connecting that pair of regions [45].

To visualizes the effects of lesions on parcels in the Lausanne anatomical atlas, we registered the lesion masks to each individual’s T1 image (the same space as the Lausanne parcel registration). We computed whether the lesion intersected > 0 voxels in that parcel, and counted the number of subjects at which that parcel was intersected by the lesion. See Fig. 1 for a visualization of the distribution of lesions across subjects and Fig. 2 for a summary of the tractography pipeline including the imputation.

**Figure 1:**
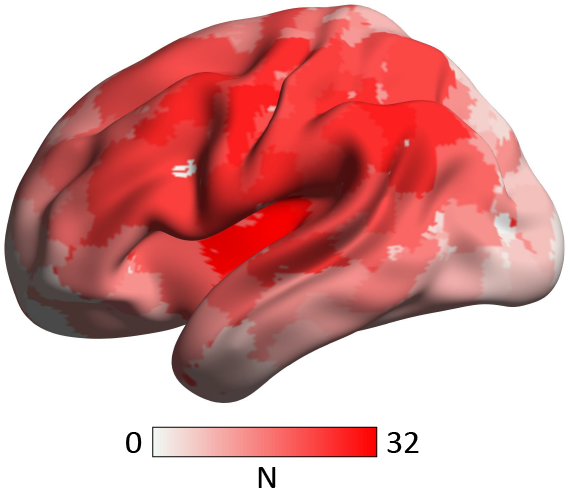
Distribution of lesioned parcels across subjects with stroke. Lesions mapped prominently to parcels in left perisylvian regions with decreasing frequency in the superior, inferior, anterior, and posterior directions. The degree of red is proportional to the number of subjects with lesions at that location. N = the number of subjects with a lesion at that parcel label.

**Figure 2:**
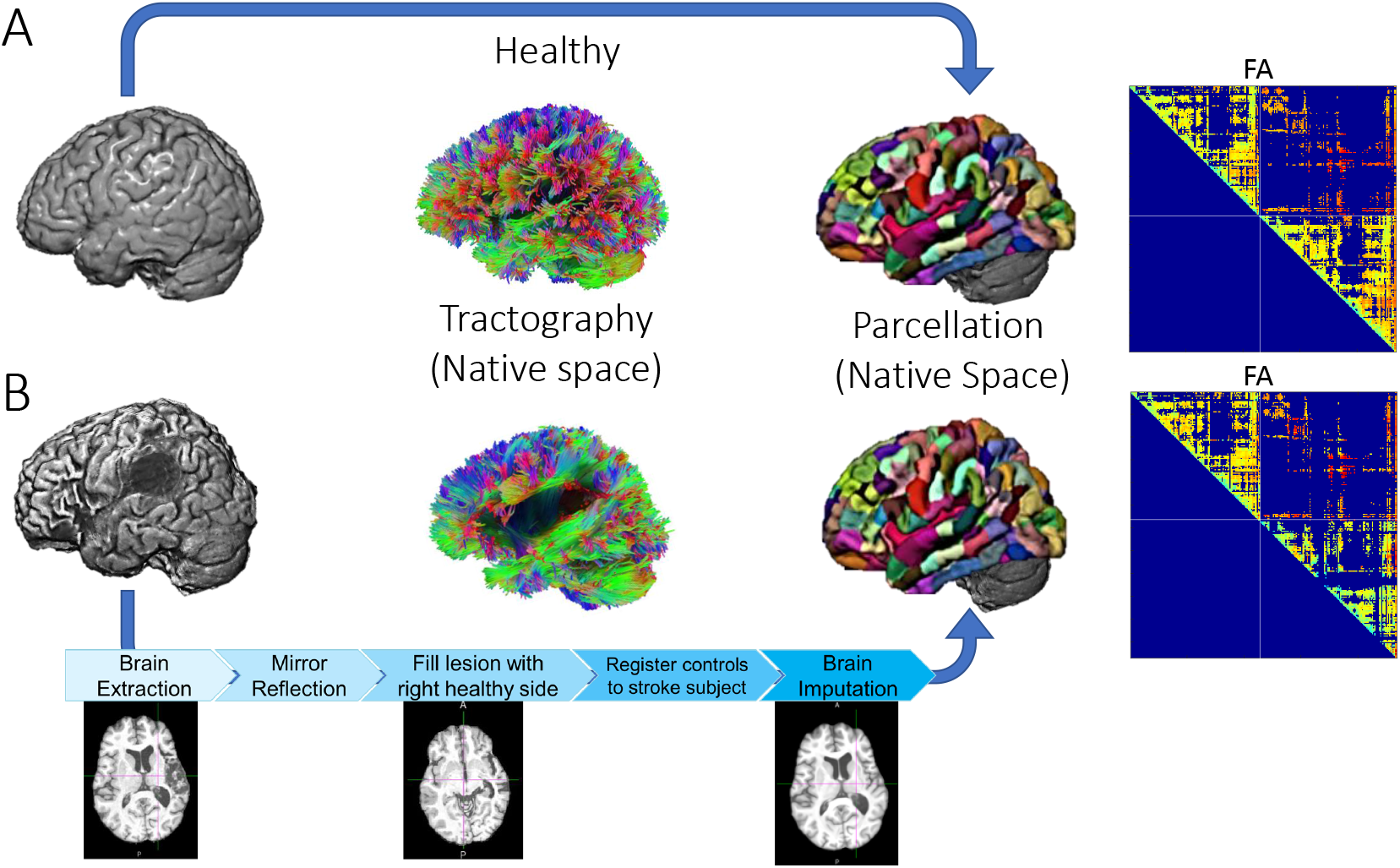
Schematic of Stroke Imputation and Diffusion Tractography. (*A*) Processing scheme for healthy subjects. Diffusion tractography was computed in subjects’ native space, and the Lausanne multiscale parcellation was fit to subjects’ anatomical T1 images. Connectomes were defined as the fractional anisotropy or streamline counts of the edges connecting each region pair and advanced to analyses. (*B*) The processing scheme for stroke subjects was the same as the healthy subjects with an additional preprocessing step. Specifically, the anatomical T1 image was imputed using the stroke subject’s right hemisphere and healthy subjects’ data to estimate the pre-lesion T1 anatomical image. The parcellation was computed on this imputed anatomical image to guide connectome extraction through the same regions as the controls.

**Figure 3:**
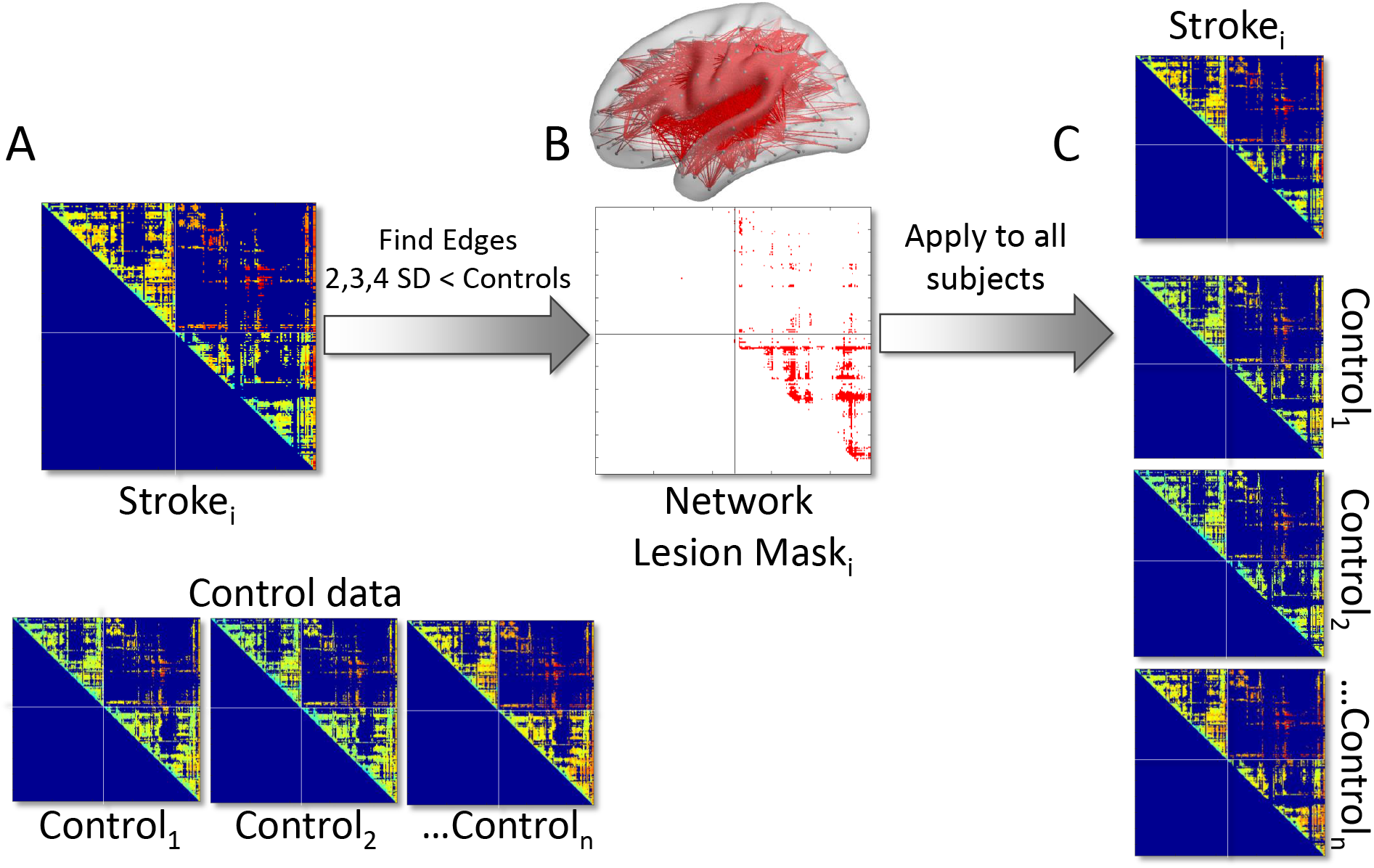
Schematic of Network Lesion Masking. (*A*), *Top* Each element *A_i_j* from each subject with aphasia (Stroke_i_) was compared to (*A*), *Bottom* the observed values in all control subjects (Control_1_ to Control_*n*_). *(B)* The elements with FA 2, 3, or 4 SDs less than controls were labeled as lesioned edges. Then, the lesion mask was applied to the stroke subject and all control subjects, and the resulting networks were advanced to connectomic analyses.

**Figure 4:**
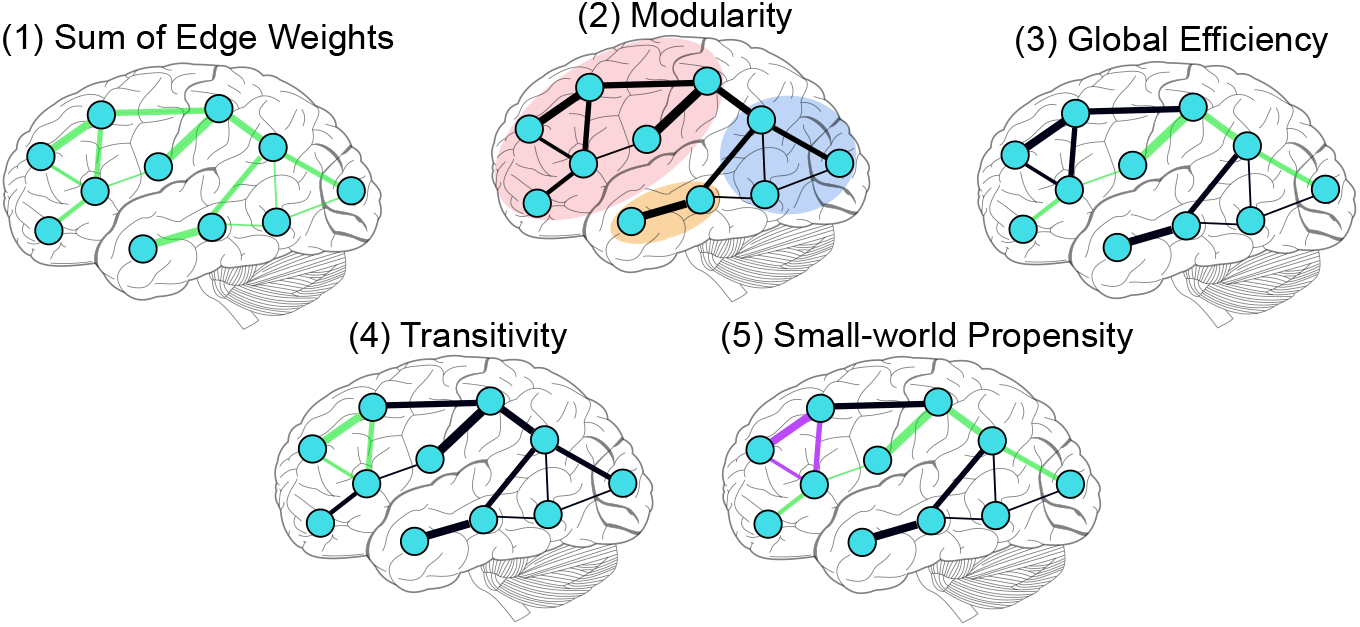
Schematic of network measures. Moving left to right from the top: we began with the (1) *average of edge weights* in each network as an overall metric capturing the density of the networks, including any edges lost due to stroke. Then, we examined the additional value of four other measures of topology compared to the sum of edge weights alone. (2) *Modularity:* measures the extent to which nodes in the network are grouped into modules (sometimes, “communities”) as a function of highly-connected nodes. (3) *Global efficiency:* one long path is represented by the set of consecutive edges highlighted in green. (4) *Transitivity:* one possible triplet’s edges are represented in green. (5) *Small-world propensity:* involves a high degree of local clustering (represented by the set of nodes connected by purple edges) and short path lengths (e.g., higher weights along the green path represents a shorter network path between the prefrontal and occipital nodes).

#### Connectome Edge Inclusion Mask

Given well-described false positives issues in diffusion tractography, it is difficult to ensure that every individual streamline is valid in the absence of ground-truth data [46]. We only permitted edges to participate in our analysis if they were present in 100% of the healthy control sample. This procedure ensured that any changes in topology observed in the stroke sample were likely to be driven by lesion-related effects rather than spurious patterns attributable to unreliable tractography findings between each pair of parcels. All triangle matrices produced via MRTrix3 were symmetrized across the diagonal prior to network analyses.

### Simulated attack

Our goal was to simulate connectome attacks using estimates of the consequences of real strokes on the connectome relative to the connectivity observed in control subjects. To identify a set of potentially lesioned edges for each subject, we compared the edge values from each stroke subject’s observed connectome to those observed in healthy subjects. Because the definition of lesioned tissue depended on a binary threshold, we computed the simulated attacks at thresholds of 2, 3, and 4 standard deviations below the mean FA or the log of the streamlines (to account for lognormal edge distributions, see [40, 47, 48, 49]) relative to the the control sample. For each threshold, a mask was created for all lesioned edges for each stroke subject. Next, we applied the edge lesion mask to each connectome in the control sample in addition to the same stroke subject. By applying the mask to the stroke subject, we ensured that the number and configuration of edges included in the analysis was equal between each stroke subject and the controls. Finally, each connectome measures was averaged across the thresholds to obtain a representative value for each subject. Then, an empirical distribution of the expected effects of lesions on network measures was obtained by computing the network measures (described below) for each possible control subject-to-lesion pairing.

### Connectome measures

We focused on five *global* network measures that are thought to characterize overall network communication. We examined (1) the network *edge weights:* the average of observed edge values, which could be related to overall network intactness and can drive other global network measures. The average edge weight served as a network proxy that estimates the overall integrity of edges in the residual connectome. We also examined the *modularity* of the networks. Modularity is thought to support local computations within tightly-connected subgroups of nodes within a network [50, 22, 51, 52] and can predict intervention-related cognitive plasticity [53]. It is computed to estimate the relative within-module connectivity of a network relative to between-module connectivity with the modularity value *Q* [54, 55].

In addition, we examined another commonly examined characteristics of the overall network topology that have been linked to variability in global cognitive performance. To examine overall network processing efficiency, we examined (2) network global efficiency [56, 57, 58, 59, 60]. Here, global efficiency is a weighted measure that quantifies average inverse shortest path length across the connectome for all pairs of nodes [61]. It is inversely related to the network measure *path length*. A pair of nodes with a short path length are connected by sequences of stronger edges. Intuitively, stronger connections between nodes can theoretically represent the strength of information flow between regions. Thus, the average path length of a network represents the extent to which all pairs of nodes are associated via short hops through the network. Accordingly, networks with high global efficiency are thought to have increased long-distance information processing capacity across all nodes mediated by short paths.

In brain networks, local clustering among u-fibers constitute most of the brain’s white matter [62]. Accordingly, short-distance, local clustering is another important aspect of information processing in brain networks. Therefore, we also examined network (3) transitivity [63, 64, 65]. Transitivity is the ratios of triangles - which are groups of three nodes connected by three non-zero edges - to all possible triplets (triangles witch edge weights equal to 1). Networks typically have many more triangles than are triplets. Therefore, greater transitivity means that there are more local clusters in a network. Networks with high transitivity are thought to have increased local communication efficiency [66].

Finally, healthy brain networks are characterized by an optimal use of available anatomical connections to support short path lengths and high clustering, which is often referred to as *small-worldness* [67, 68]. Small-world brain networks are thought to confer many of the processing advantages that support diverse and dynamic cognitive functions [30]. To investigate this property, we used a robust measure of small-worldness, (4) *small-world propensity* (SWP) [69]. SWP is a weighted metric for small-worldness that accounts for networks of different densities, standardizing the measure against individualized network null models. This technique makes SWP appropriate for measuring small-worldness in weighted networks by mitigating the network density-dependence of other measures.

### Statistical analyses

#### The effects of lesions on connectome measures and behavior

First, we examined whether observed and simulated strokes had significant effects on each network measure in the whole bran and within the left hemisphere. We computed Welch’s t-tests assuming unequal variances using Satterthwaite’s approximation for degrees of freedom for each measure against those observed in the control subjects corrected for multiple comparisons at an alpha level of 0.05. Then, we used bootstrapping to estimate the proportion of network measure sample means from the simulated attacks that fell within the range of the observed lesion for each measure. We used this technique because we intended the simulation to sample from all lesion-control subject pairs to yield a distribution of possible lesion profiles in a much larger simulated sample. Specifically, we performed 10,000 resamplings with replacement of 40 subjects and quantified the proportion of simulated attack sample means for each network measure (FA and streamline) and each size network (whole brain or left hemisphere). Finally, to estimate the behavioral variance accounted for by lesion volumes, we fit separate linear regression models using lesion volume as an independent variable and either WAB-AQ, the factor sum score, or each behavioral factor score corrected for multiple comparisons at an alpha level of 0.05.

#### Preparing network measures to identify behavioral variance beyond lesion volume

Our objective was to obtain and present an empirical estimate for the full range of possible lesion-behavior relationships observed in the real, simulated, and null analyses.

For all connectome analyses, we were interested in the total variance accounted for using all five network measures for the observed and simulated data. Prior to analyses, we tested the network measures for violations of normality with the Kolmogorov-Smirnov test. To correct for skewed distributions in the observed statistics, we used a log-transformation for global efficiency and SWP. In the simulated attack statistics, we observed negative values and skew for each statistic; thus, we added a constant value of 1 to each measure prior to a log-transformation. Finally, network measures were standardized using z-scores prior to all analyses.

To test the hypothesis that real and simulated measures of network topology were related behavioral scores beyond lesion volume, we first computed separate linear regressions with using stroke subject’s lesion volume as the independent variable and each behavioral scale as the dependent variable. Then, we used the residualized behavioral scores as the dependent variable for all connectome-behavior analyses. We performed the same procedure for each network measure to mitigate any remaining influences of lesion volume effects.

#### Network-behavior relationships in observed, simulated, and null regression models

Next, all analyses associating network measures with behavioral scores were performed using linear regression in R statistical software [70]. Our simulated attacks broke the relationship between stroke subjects and behavior by randomly sampling residual anatomical connectomes after simulating strokes in control subjects, but preserved the relationship between each behavioral score and simulated lesion. The null model completely randomized the relationships between simulated lesions and behaviors.

Specifically, we computed linear regression models for (1) the observed stroke network topology, (2) the simulated stroke network topology (10,000 permutations per lesion edge threshold), and (3) a randomized shuffling of all simulated network measures against the behavior (10,000 permutations per lesion edge threshold). We examined the effects of observed and simulated lesions on each of the topological measures in the whole brain and within intra-left hemisphere connectomes (i.e., only the whole-brain connectomes included interhemispheric and right-hemispheric fibers). Then, we computed the relationships between network measures and the behavioral scores for the whole brain and intra-left hemisphere connectomes.

In analyses of the observed data (i.e., data from subjects with real strokes), we used each of the five network measures as independent variables (z-scored across subjects) and each behavioral score as a dependent variable (raw WAB scores, the factor sum scores, or one of the four behavioral factor scores) in separate linear regression models. Because the network measures and behavioral scores were the residuals obtained after regressing out the influence of lesion volume, we obtained specific parameter estimates for each network measure and the total variance accounted for in the models (R^2^ value) beyond lesion volume.

To obtain estimates for network-behavior relationships in the simulated attack, we computed the linear regression models with the same dependent behavioral variables, but with independent network variables sampled from the control-lesion pairings for the simulated attack (z-scored across subjects). Specifically, in 10,000 permutations, we randomly assigned each stroke lesion to a healthy brain from the control sample while preserving the link between that lesion and the behavioral outcome. We then computed each network measure on that sample of simulated attacks, and fit a regression model. Across the 10,000 models, this approach provided a full representation of the absolute minimum, maximum, and of the predictive value (*R*^2^) of the anatomical connectomes beyond lesion volume. In addition, we obtained the range of beta weights for each network measure in the simulations to reveal their relative contributions to the prediction.

In high-dimensional data analyses such as these, it is often helpful to have an empirical null distribution to contextualize the models of interest. By fully randomizing the relationships between the simulated and network measures and behavior (i.e., breaking the lesion-behavior pairing in the simulated strokes), we were able to create a distribution of the expected variance accounted for (*R*^2^) if the data were to be completely randomized. In this kind of null permutation analysis, we would expect that models could trivially account for more variance than 0 by chance. Further, the null could include permutations equaling or similar to real lesion-behavior network pairings, potentially meeting or exceeding the variances obtained from the observed or simulated attack. If the observed or simulated models exhibited effects that were higher that the null’s central tendency, it would increase our confidence that the network topology across the range of simulated attack outcomes is non-trivially related to behavior above and beyond lesion volumes.

Our primary goal was to test whether the simulated attacks based on observed lesions differed from the null distribution. After computing the *R*^2^ value for each null and simulated model in each permutation, we used a 2 (simulated *versus* null) x 6 (behavioral variables) ANOVA to test the effects of (1) the simulated attack relative to the null permutations and (2) behavioral domain on the estimated *R*^2^ values.

## Results

### The effects of lesions on network measures

The observed strokes influenced network topology for each measure. In FA networks in the whole brain, we observed a reduction in mean edge weight, network efficiency, and increased SWP. In the left hemisphere, we observed all of these effects in addition to reduced modularity (see Fig. 5). In the streamline networks in the whole brain, we observed reduced mean edge weight, network efficiency, and transitivity as well as increased modularity. In the left hemisphere, we observed reduced edge weights, modularity and increased transitivity (see Fig. 5, Table 2). The network measures from the simulated strokes were similar to those observed after real strokes (see Table 3), suggesting that they were reasonable approximations of stroke effects on the connectomes.

**Figure 5:**
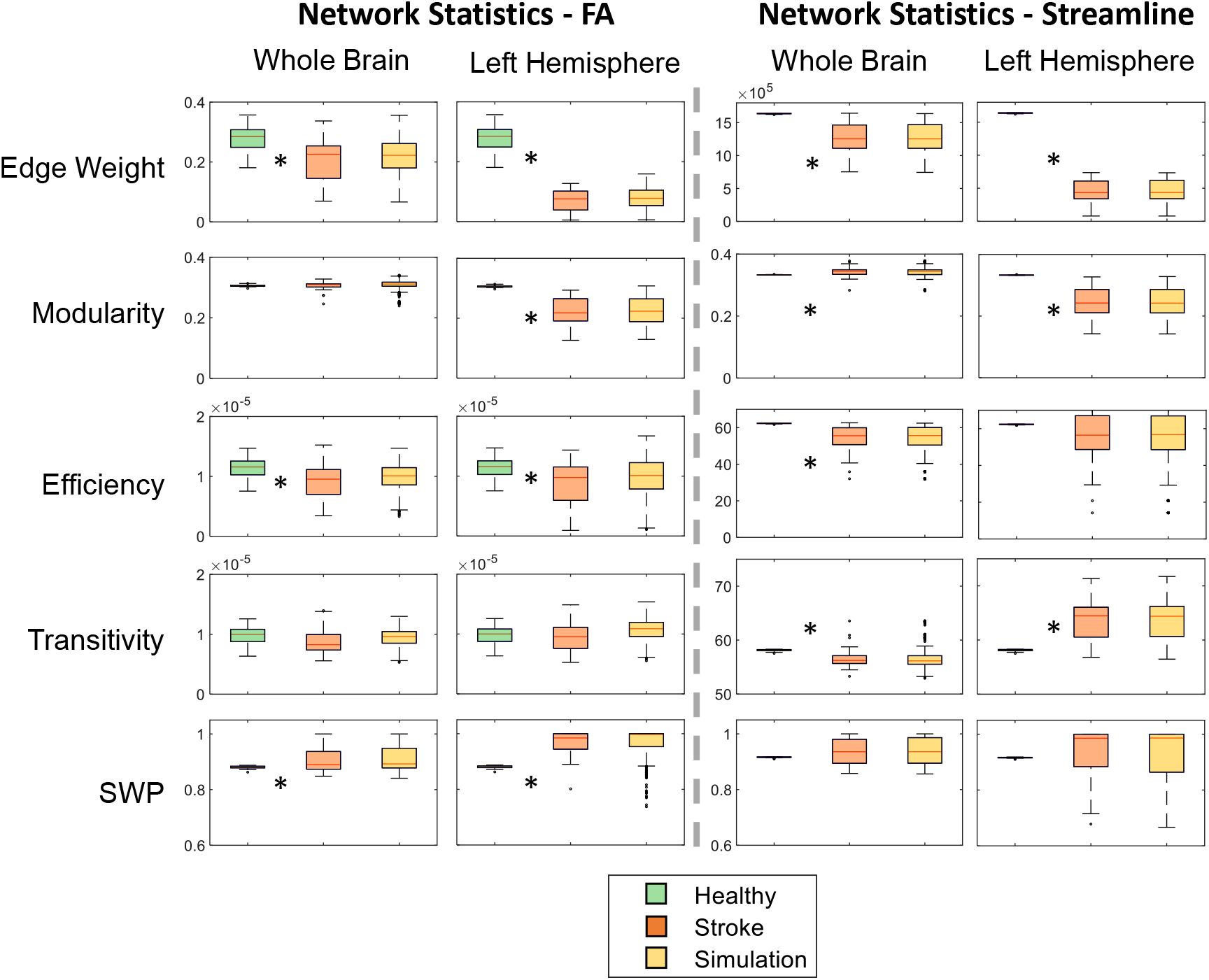
The effects of stroke on network measures in observed and simulated attack connectomes. The leftmost column of each plot facet shows the network statistic observed in controls, followed by that observed in strokes, then the simulated attacks. Network measures are presented in their raw (untransformed) values prior to inclusion in network-behavior analyses. Asterisks indicate a significant Welch’s two-sample t-test between the control and stroke network measures at *p* < 0.001 (a stringent threshold after Bonferroni correction for 40 total tests in FA and Streamline data). The top and bottom edges of the boxes represent the 25th and 75th percentiles, respectively. SWP = small world propensity.

**Table 2:**
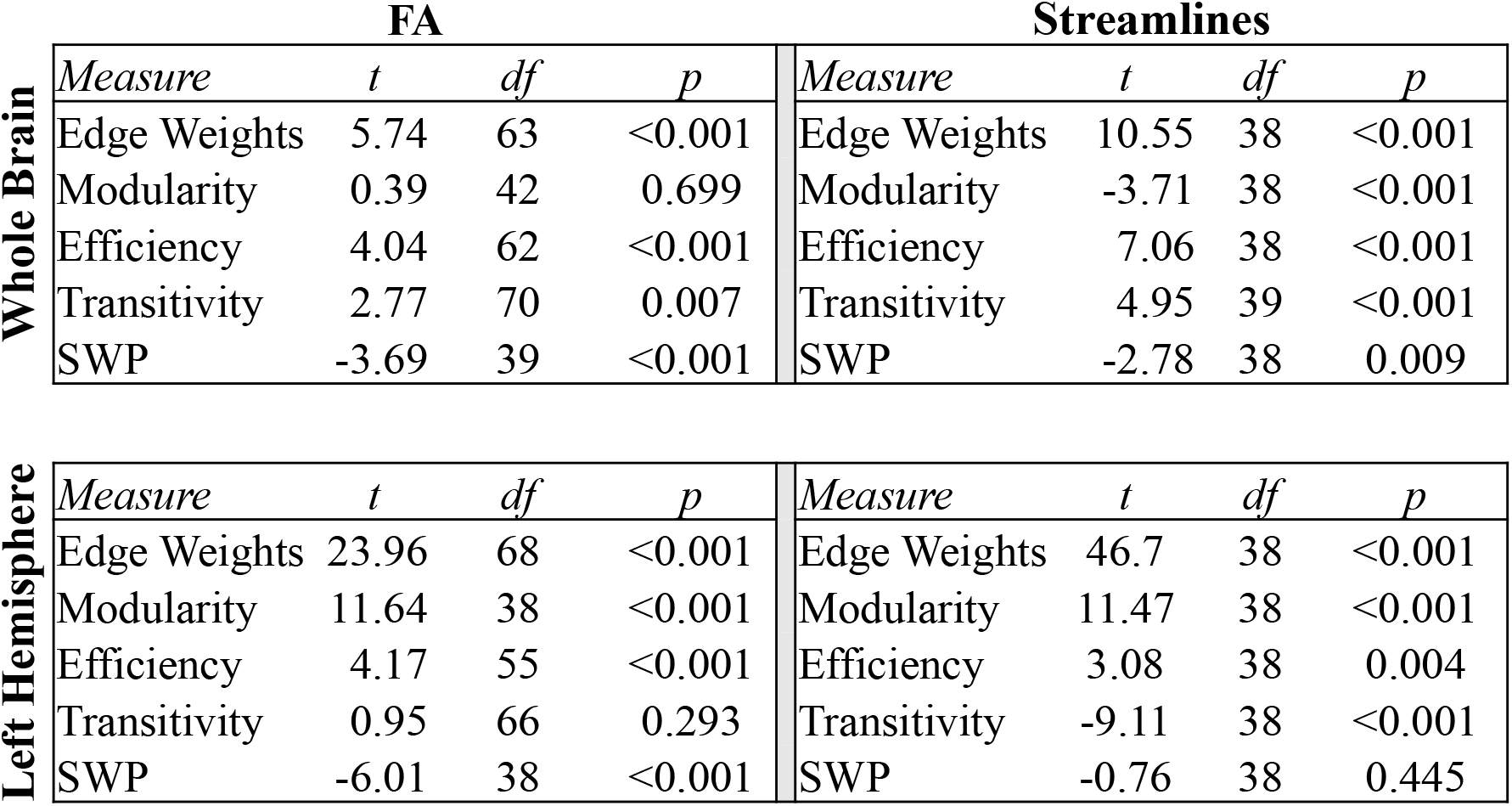
Two-samples t-tests assuming unequal variances comparing network measures between the control and stroke samples for the whole brain FA networks (df = Satterthwaite’s approximation for samples with unequal variances).

|  | FA |  |  |  | Streamlines |  |  |  |
| --- | --- | --- | --- | --- | --- | --- | --- | --- |
|  | *Measure* | *t* | *df* | *p* | *Measure* | *t* | *df* | *p* |
| Whole Brain | Edge Weights | 5.74 | 63 | <0.001 | Edge Weights | 10.55 | 38 | <0.001 |
|  | Modularity | 0.39 | 42 | 0.699 | Modularity | -3.71 | 38 | <0.001 |
|  | Efficiency | 4.04 | 62 | <0.001 | Efficiency | 7.06 | 38 | <0.001 |
|  | Transitivity | 2.77 | 70 | 0.007 | Transitivity | 4.95 | 39 | <0.001 |
|  | SWP | -3.69 | 39 | <0.001 | SWP | -2.78 | 38 | 0.009 |
| Left Hemisphere | Edge Weights | 23.96 | 68 | <0.001 | Edge Weights | 46.7 | 38 | <0.001 |
|  | Modularity | 11.64 | 38 | <0.001 | Modularity | 11.47 | 38 | <0.001 |
|  | Efficiency | 4.17 | 55 | <0.001 | Efficiency | 3.08 | 38 | 0.004 |
|  | Transitivity | 0.95 | 66 | 0.293 | Transitivity | -9.11 | 38 | <0.001 |
|  | SWP | -6.01 | 38 | <0.001 | SWP | -0.76 | 38 | 0.445 |

**Table 3:** The proportion of bootstrap sample means of simulated attacks compared the range of observed lesion values.

| Condition | Edge Weight | Modularity | Efficiency | Transitivity | SWP |
| --- | --- | --- | --- | --- | --- |
| FA - Whole Brain | 1.00 | 1.00 | 1.00 | 1.00 | 1.00 |
| FA - Left Hemisphere | 1.00 | 1.00 | 1.00 | 1.00 | 0.77 |
| Streamline - Whole Brain | 1.00 | 1.00 | 1.00 | 1.00 | 1.00 |
| Streamline - Left Hemisphere | 1.00 | 1.00 | 1.00 | 1.00 | 0.77 |

### Relationships between observed and simulated connectome topology and aphasia-related behaviors

Lesion volume accounted for approximately 44% of WAB-AQ, 53% of the factor sum, and 10-16% of the variance in factor scores (Table 4). In addition, lesion volume was negatively correlated with FA network edge weights, modularity, and efficiency, and positively associated with small world propensity. Lesion volume was negatively correlated with streamline network edge weights, efficiency, and positively associated with transitivity and small world propensity (Table 5).

**Table 4:**
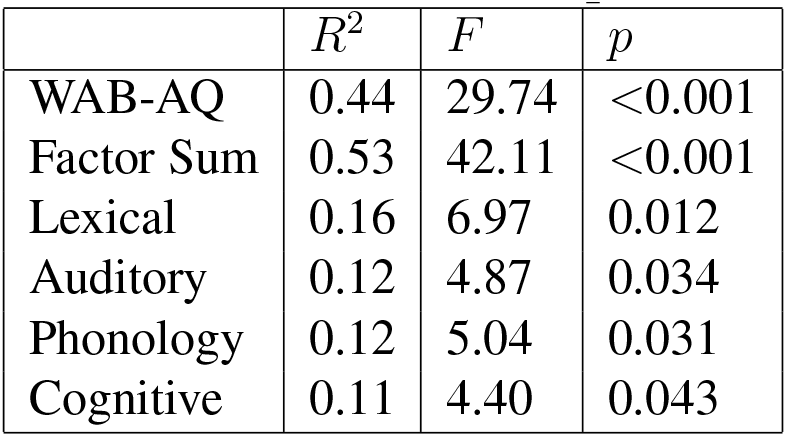
The relationships between lesion volume and behavioral scores.

**Table 5:** The relationships between lesion volume and network measures. R-values represent Pearson’s correlation coefficients between lesion volume and the network measure.

|  | *Measure* | *R* | *p* |
| --- | --- | --- | --- |
| **FA** | Edge Weights | -0.52 | 0.001 |
|  | Modularity | -0.44 | 0.005 |
|  | Efficiency | -0.46 | 0.003 |
|  | Transitivity | -0.13 | 0.414 |
|  | SWP | 0.34 | 0.033 |
| **Streamline** | Edge Weights | -0.87 | 0.000 |
|  | Modularity | -0.21 | 0.201 |
|  | Efficiency | -0.89 | 0.000 |
|  | Transitivity | 0.55 | 0.000 |
|  | SWP | 0.31 | 0.051 |

We regressed lesion volume from the behavioral and network data to obtain the additional unique variance between the network measures and behavior. The full model results are presented in Fig. 6. For each of the FA and streamline whole brain and intra-left hemisphere networks, the variance accounted for (*R*^2^) by the simulated attack networks was greater than the nulls in the omnibus ANOVA (see Table 6 for condition-wise marginal means, which quantify the difference in the observed *versus* the null (*R*^2^) across behaviors).

**Figure 6:**
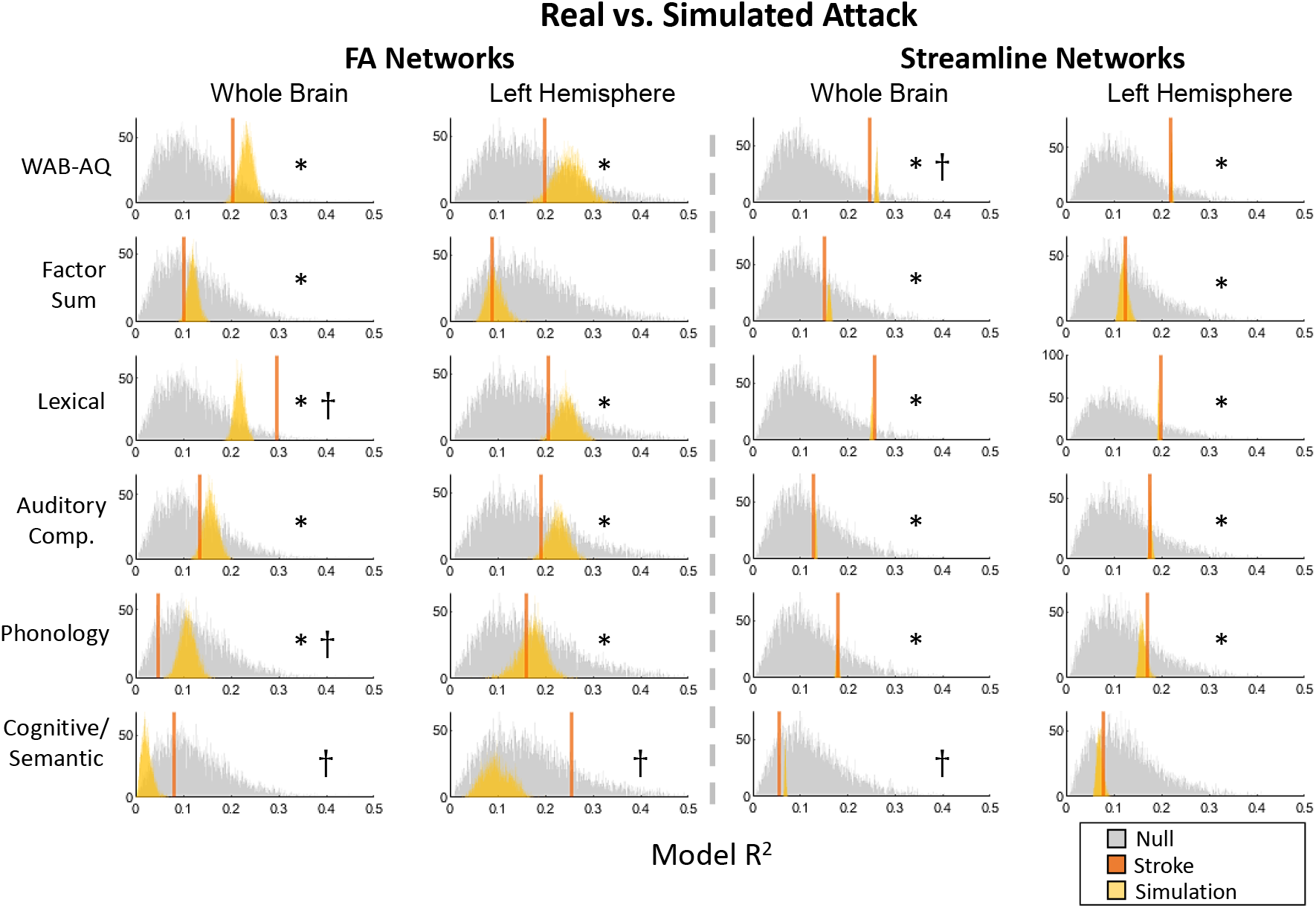
Network measures and behavioral variance in FA networks. Each plot facet illustrates histograms of the simulated null, histograms of the simulated attack distributions, and the observed *R*^2^ with solid vertical lines. Asterisks indicate significant *post hoc* Welch’s one-tailed t-tests assuming unequal variances comparing the *R*^2^ values in the simulated attacks to the null distribution at p < 0.001. Daggers indicate cases where the observed *R*^2^ value was outside the range obtained in the simulated attack models.

**Table 6:** Marginal means for the simulated attack relative to the null for FA and streamline networks in the whole brain and left hemisphere.

| Type | Location | Lower CI | Mean Diff. | Upper CI | p |
| --- | --- | --- | --- | --- | --- |
| FA | Whole | 0.017 | 0.017 | 0.016 | <0.001 |
| FA | Left | 0.01 | 0.009 | 0.008 | <0.001 |
| Streamline | Whole | 0.048 | 0.047 | 0.047 | <0.001 |
| Streamline | Left | 0.03 | 0.03 | 0.029 | <0.001 |

In addition, there were main effects of behavior on the model *R*^2^ values (see Supplementary Tables 1–4). The central tendency of the null models revealed that network measures would be expected to account for nearly 10% of the behavioral variance on global or specific language performance at random (i.e., when the link between the lesion being simulated and behavioral score was broken). Outperforming the null, the simulated attack models generally suggested that about 20% of global aphasia outcomes measured with the WAB-AQ could be accounted for residual anatomical network topology. Among the language behavioral factor scores, lexical processing exhibited the strongest relationships with network topology, at 20% or more variance accounted for by the whole brain or left hemisphere network measures. In most cases, the observed *R*^2^ estimate was within the range estimated in the simulated attacks. Exceptions were observed in several cases, and more frequently in FA networks, where observed stroke estimates were outside the simulated estimates for the whole-brain lexical, phonology, and cognitive/semantic factors and the left-hemisphere cognitive/semantic factors.

### Beta weights for specific network measures

We additionally obtained the beta values for the simulated attack models to observe which measures contributed the most weight to model R^2^. Beta weights for the whole brain and intra-left hemisphere models are illustrated in Figures 7. In the whole brain models, edge weights and efficiency were most consistently associated with higher betas across behaviors, with some variation across the individual factor scores. Within the left hemisphere, edge weights and efficiency remained relatively stronger contributors to the global WAB and factor score sum behavioral measures. Among the four factor subscores, network measure beta weights exhibited more variation across specific factors.

**Figure 7:**
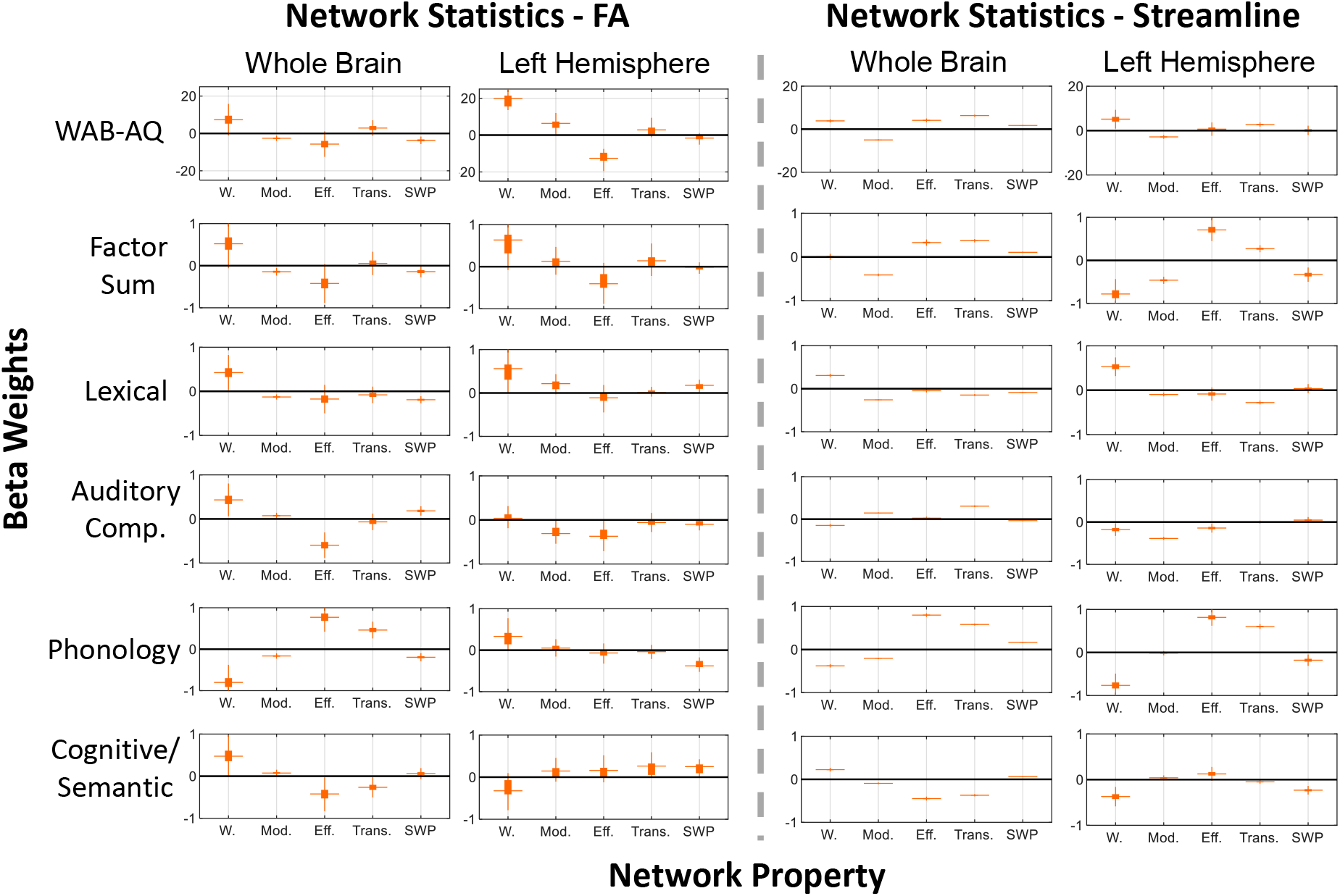
Simulated attack estimated beta weights for each network measure in FA networks. Each plot represents the range of betas obtained from the simulated attack models for the network measure. The top and bottom edges of the boxes represent the 25th and 75th percentiles, respectively. W. = edge weights, Mod. = modularity, Eff. = Efficiency, Trans. = Transitivity, SWP = small world propensity.

See Supplementary Table 1 for observed and simulated model betas.

## Discussion

In subjects with left-hemispheric strokes, we used an anatomical network simulated attack analysis to examine the relationships between topological network measures, a widely used clinical aphasia outcome measure (the WAB-AQ), and dimensional factor scores of language performance. We found that (1) simulating lesions can provide a good approximation of observed lesion effects on networks that do not suggest widespread network plasticity across people, (2) that the network properties of the simulated lesions can explain variance in behavior above and beyond lesion size, (3) that in most cases observed lesions do not explain more variance in behavior than simulated lesions, and (4) that relationships of simulated and observed lesions to behavior differ depending on the behavior being examined.

In general, lesions in the left hemisphere disrupted several measures of global topological organization in persons with aphasia post-stroke. In whole brain and left hemispheric FA networks, stroke reduced the overall edge weight and efficiency relative to controls. In contrast, small-world propensity tended to increase post-stroke, which recapitulates the reduced network efficiency compared to relatively consistent measure of global clustering (transitivity) [69]. Interestingly, modularity was decreased in the intra-hemispheric connectomes. The relative increases in modularity in the whole brain were likely driven by the absorption of residual left-hemispheric networks into right hemispheric homotopic communities via interhemispheric fibers mediated through the corpus callosum. These patterns were similar in the simulated attacks, suggesting that the simulations were reasonable approximations of real stroke effects. A minority of sampled simulated cases in the left hemisphere had SWP values averaging outside the range of stroke subjects, potentially reflecting neuroplastic or premorbid differences in the hemispheric balance between long distance and local communication.

As a topic of focus in several previous studies, modularity changes in persons with stroke could vary based on the location of lesions. For instance, left hemisphere anatomical modularity has been found to increase in subjects with upper limb motor deficits [71], and increased left hemisphere anatomical modularity have been associated with more severe chronic aphasia [22]. In contrast, reduced modularity in functional connectomes observed in multiple stroke phenotypes [72] has been shown to partially recover in the transition from the acute to chronic phase [73, 23, 52]. Anatomical connections and network topology predict region-to-region functional connectivity [74], and it will be important to clarify how specific lesion distributions interact with anatomy, and joint anatomy-function relationships. For instance, sensorimotor cortices are highly interconnected within each hemisphere, and precentral regions are often revealed to participate in the brain’s anatomical hub system [75, 76]. Thus, disrupting sensorimotor regions and their connections is likely to enhance the modularity of the remaining intra-hemispheric network. Reduced modularity was not as strongly or consistently related to the language behaviors examined here, suggesting that these topological changes might not be as uniquely informative as other measures of topology (e.g., especially the overall connections in the network). Overall, the topology of the left hemisphere was sparser and more tightly clustered, which is often thought to limit the general ability for a network to transmit information and reduce the interference between competing demands on the network due to the loss of specialized processing modules [68].

In the simulated attack regression models associating brain network measures with behavior, we quantified the variance accounted for by anatomical network measures above and beyond the effects of lesion volume. Unsurprisingly, lesion volume accounted for a moderate amount of variance in the global aphasia measures (WAB-AQ and factor sum), and less variance in the specific factor scores. The network measures and behavior residualized for lesion volume revealed that additional variance was accounted for by anatomical network topology. Among the network-behavior analyses, in most cases, the observed model value was within the simulated attack distribution, placing the observed *R*^2^ value within a few percent of the central tendency of the simulated distribution. We did not obtain evidence that the observed brain-behavior relationships always significantly over- or under-perform estimates from simulated attacks for the measures we examined. These findings suggest that the majority of brain-behavior relationships in the simulated permutations are driven by the direct effects of the lesion on the connectome - i.e., that the simulated attacks were an informative basis to obtain confidence intervals for the effects of prototypical lesions on the connectome and behavior. Cases where specific deviations between the observed and simulated models were observed (e.g., in the FA lexical, phonological, and semantic factors) could reveal the influences of sampling effects, premorbid network organization and behavior, deafferentiation, diaschisis, adaptive neuroplasticity, or related neurological effects [77].

Across the simulations, there was an intuitive relationship between measures of global network topology and overall aphasia severity, accounting for 20 % of the variance on the WAB-AQ. Interestingly, this network-behavior relationship was stronger than that observed with the total factor sum score. Within the left hemisphere, this pattern remained, and lexical processing and auditory comprehension tended to have strong relationships with network topology, potentially reflecting the anatomically distributed demands of these tasks in left hemispheric perisylvian circuits [78], association regions [79], and sensory-perceptual pathways [80, 81]. Perhaps due to the relatively circumscribed circuits thought to mediate phonological processing [82, 83] and relatively preserved prefrontal circuitry that might mediate the functions in our cognitive factor [84, 85], we observed weaker relationships between these behaviors and the topology in the simulated attacks.

Overall, streamline networks appeared to offer substantially more reliable point estimates for global and dimensional behavioral outcomes despite similar central tendencies to FA networks. This could be because our use of a reliable healthy connectome ensured consistent sets of streamline edges in healthy controls, whereas FA values are derived from estimated streamlines and offer an additional source of variance in the simulations. In addition, streamline distributions tend to be heavy-tailed with few highly connected pairs of regions [40, 47, 48, 49] with node degree (number of connections) and strength (the total weight of the connections) distributions that follow exponentially-truncated power laws [86, 87]. Lesions induce a significant loss in the number of estimated fibers, and the measured topology of these losses across subjects within the reliable healthy connectome will be strongly influenced by the heavy-tailed distribution of edges and presence of a subset of high-connection-strength nodes. In contrast, FA is computed over the estimated streamlines connecting regions pairs regardless of their number, exhibiting significantly less skew and consequently relatively fewer high-connection-strength nodes. Qualitatively, FA can represent the integrity of axonal pathways [88, 89], offering a distinct interpretive value relative to streamlines. However, our results suggest that increased caution when evaluating the effect sizes of brain-behavior relationships could be advised for FA relative to streamline connectomes.

Across the FA networks, it was not clear that any one of the investigated network measures uniquely corresponds to a single dimension of language function post-stroke. When examined using the whole brain FA networks, edge weights and network global efficiency tended to contribute to most of the behavioral measures. This finding could represent the possibility that individual variation in the myriad nearby, short-distance connections such as u-fibers and direct connections are relevant to mediating recovery [20, 90]. U-fibers dominate the brain’s white matter but their links to cognition are conspicuously understudied [91]. Intuitively, a higher degree of intact local bypasses could facilitate adaptations to lost functions in general [92]. It is likely that as the behavioral measures increase in specificity, the unique edges contributing to losses in each function vary, driving differences in topology-behavior relationships [14]. In lexical processing (where the models accounted for the most variance among the individual factor scores) edge weights were most prominently related to behavior for both streamlines and FA, suggesting that overall loss of connections independent of their relationship with lesion volume is a key mediator of deficits relative to other topological measures.

Several limitations to our work can motivate future studies. We focused on a narrow set of commonly used network topology measures that characterize some of the aspects of global network organization as an initial benchmark for the connectome bases of language performance. Numerous other measures are available, but the link between specific network measures and cognitive functions remains an active area of inquiry and debate [6, 93, 94, 95]. More specific hypotheses that allow researchers to rule out spurious or non-specific network-behavior effects, ideally informed by theoretical models, should be a focus of applied network studies. In addition, other measures that represent connectome edges (streamline density, diffusivity, etc., [96, 34, 41]) could be investigated. We recommend that these efforts will be best supported by collaborative efforts to pool patient samples and test the robustness of brain-behavior relationships. In addition, we used a well-established anatomical atlas to guide our parcellation with an imputation procedure to compare healthy and stroke subjects, but numerous atlases are now available. Given that there is no consensus that a particular atlas is ideal for any specific purpose [97], we further encourage researchers to collaboratively pool data to examine the reliability and validity of different processing decisions in anatomical and functional studies [98, 99, 100, 101, 102, 103].

## Conclusion

Our simulations revealed that several anatomical connectome measures thought to be related to global network processing can be expected to account for 10-20% of the variance in language performance on clinical measures above and beyond lesion volume. Importantly, measures of whole brain and left hemisphere anatomical connectomes have stronger relationships with global language function, reflecting an intuitive relationship between network-wide integrity and overall functioning. More specific measures of anatomical circuits could be necessary to gain more sensitivity to distinct language processes. Simulated attacks are useful in leveraging matched comparison samples to obtain confidence estimates for observed effects. Differences between observed and simulated values could identify the influences of premorbid status, deafferentation, diaschisis, and neuroplasticity following stroke.

## Acknowledgments

JDM acknowledges support from NIH grants DP5-OD-021352, R01-DC-16800, R01-AG-059763, and Department of the Army grant PRMRP 12902164. PET and JDM acknowledge support from NIH grant R01-DC014960.

---

**Supplementary Table 1:** FA marginal means for behavior in the whole brain.

| Behavior 1 | Behavior 2 | Lower CI | Mean Diff. | Upper CI | p |
| --- | --- | --- | --- | --- | --- |
| WAB | Factor Sum | 0.055 | 0.056 | 0.058 | <0.001 |
| WAB | Lexical | 0.006 | 0.007 | 0.009 | <0.001 |
| WAB | Auditory Comp. | 0.036 | 0.038 | 0.039 | <0.001 |
| WAB | Phonology | 0.06 | 0.062 | 0.063 | <0.001 |
| WAB | Cognitive/Semantic | 0.103 | 0.104 | 0.106 | <0.001 |
| Factor Sum | Lexical | -0.05 | -0.049 | -0.047 | <0.001 |
| Factor Sum | Auditory Comp. | -0.02 | -0.019 | -0.017 | <0.001 |
| Factor Sum | Phonology | 0.004 | 0.005 | 0.007 | <0.001 |
| Factor Sum | Cognitive/Semantic | 0.047 | 0.048 | 0.05 | <0.001 |
| Lexical | Auditory Comp. | 0.029 | 0.03 | 0.032 | <0.001 |
| Lexical | Phonology | 0.053 | 0.054 | 0.056 | <0.001 |
| Lexical | Cognitive/Semantic | 0.096 | 0.097 | 0.099 | <0.001 |
| Auditory Comp. | Phonology | 0.023 | 0.024 | 0.026 | <0.001 |
| Auditory Comp. | Cognitive/Semantic | 0.065 | 0.067 | 0.068 | <0.001 |
| Phonology | Cognitive/Semantic | 0.041 | 0.043 | 0.044 | <0.001 |

**Supplementary Table 2:** FA marginal means for behavior in the left hemisphere.

| Behavior 1 | Behavior 2 | Lower CI | Mean Diff. | Upper CI | p |
| --- | --- | --- | --- | --- | --- |
| WAB | Factor Sum | 0.075 | 0.078 | 0.08 | <0.001 |
| WAB | Lexical | 0.001 | 0.003 | 0.005 | 0.004 |
| WAB | Auditory Comp. | 0.01 | 0.012 | 0.014 | <0.001 |
| WAB | Phonology | 0.037 | 0.039 | 0.042 | <0.001 |
| WAB | Cognitive/Semantic | 0.074 | 0.077 | 0.079 | <0.001 |
| Factor Sum | Lexical | -0.077 | -0.075 | -0.072 | <0.001 |
| Factor Sum | Auditory Comp. | -0.068 | -0.066 | -0.064 | <0.001 |
| Factor Sum | Phonology | -0.041 | -0.039 | -0.036 | <0.001 |
| Factor Sum | Cognitive/Semantic | -0.004 | -0.001 | 0.001 | 0.632 |
| Lexical | Auditory Comp. | 0.007 | 0.009 | 0.011 | <0.001 |
| Lexical | Phonology | 0.034 | 0.036 | 0.039 | <0.001 |
| Lexical | Cognitive/Semantic | 0.071 | 0.074 | 0.076 | <0.001 |
| Auditory Comp. | Phonology | 0.025 | 0.027 | 0.03 | <0.001 |
| Auditory Comp. | Cognitive/Semantic | 0.062 | 0.065 | 0.067 | <0.001 |
| Phonology | Cognitive/Semantic | 0.035 | 0.037 | 0.04 | <0.001 |

**Supplementary Table 3:** Streamline marginal means for behavior in the whole brain.

| Behavior 1 | Behavior 2 | Lower CI | Mean Diff. | Upper CI | p |
| --- | --- | --- | --- | --- | --- |
| WAB | Factor Sum | 0.049 | 0.05 | 0.052 | <0.001 |
| WAB | Lexical | 0.003 | 0.005 | 0.006 | <0.001 |
| WAB | Auditory Comp. | 0.063 | 0.064 | 0.066 | <0.001 |
| WAB | Phonology | 0.04 | 0.041 | 0.043 | <0.001 |
| WAB | Cognitive/Semantic | 0.095 | 0.096 | 0.098 | <0.001 |
| Factor Sum | Lexical | -0.047 | -0.045 | -0.044 | <0.001 |
| Factor Sum | Auditory Comp. | 0.013 | 0.014 | 0.016 | <0.001 |
| Factor Sum | Phonology | -0.01 | -0.009 | -0.007 | <0.001 |
| Factor Sum | Cognitive/Semantic | 0.045 | 0.046 | 0.048 | <0.001 |
| Lexical | Auditory Comp. | 0.058 | 0.059 | 0.061 | <0.001 |
| Lexical | Phonology | 0.035 | 0.036 | 0.038 | <0.001 |
| Lexical | Cognitive/Semantic | 0.09 | 0.091 | 0.093 | <0.001 |
| Auditory Comp. | Phonology | -0.025 | -0.023 | -0.022 | <0.001 |
| Auditory Comp. | Cognitive/Semantic | 0.031 | 0.032 | 0.034 | <0.001 |
| Phonology | Cognitive/Semantic | 0.054 | 0.055 | 0.057 | <0.001 |

**Supplementary Table 4:** Streamline marginal means for behavior in the whole brain.

| Behavior 1 | Behavior 2 | Lower CI | Mean Diff. | Upper CI | p |
| --- | --- | --- | --- | --- | --- |
| WAB | Factor Sum | 0.049 | 0.05 | 0.052 | <0.001 |
| WAB | Lexical | 0.012 | 0.014 | 0.015 | <0.001 |
| WAB | Auditory Comp. | 0.021 | 0.022 | 0.024 | <0.001 |
| WAB | Phonology | 0.029 | 0.03 | 0.032 | <0.001 |
| WAB | Cognitive/Semantic | 0.074 | 0.075 | 0.077 | <0.001 |
| Factor Sum | Lexical | -0.038 | -0.036 | -0.035 | <0.001 |
| Factor Sum | Auditory Comp. | -0.029 | -0.028 | -0.026 | <0.001 |
| Factor Sum | Phonology | -0.021 | -0.02 | -0.018 | <0.001 |
| Factor Sum | Cognitive/Semantic | 0.024 | 0.025 | 0.027 | <0.001 |
| Lexical | Auditory Comp. | 0.007 | 0.008 | 0.01 | <0.001 |
| Lexical | Phonology | 0.015 | 0.016 | 0.018 | <0.001 |
| Lexical | Cognitive/Semantic | 0.06 | 0.062 | 0.063 | <0.001 |
| Auditory Comp. | Phonology | 0.006 | 0.008 | 0.01 | <0.001 |
| Auditory Comp. | Cognitive/Semantic | 0.052 | 0.053 | 0.055 | <0.001 |
| Phonology | Cognitive/Semantic | 0.044 | 0.045 | 0.047 | <0.001 |

**Supplementary Table 5:** Beta weights for each condition of the observed and simulated regression models.

FA - Whole Brain
| Behavior | Edge Weights | Modularity | Efficiency | Transitivity | SWP |
| --- | --- | --- | --- | --- | --- |
| WAB-AQ | 35.14 | -0.66 | -21.81 | -12.76 | -3.37 |
| Factor Sum | 1.93 | 0.00 | -1.34 | -0.46 | -0.07 |
| Lexical | 1.40 | -0.05 | -0.04 | -1.34 | -0.24 |
| Auditory Comp. | 0.20 | 0.18 | -0.90 | 0.44 | 0.04 |
| Phonology | 0.72 | -0.13 | -0.33 | -0.26 | 0.09 |
| Cognitive/Semantic | -0.38 | -0.01 | -0.07 | 0.70 | 0.04 |

FA - Left Hemisphere
| Behavior | Edge Weights | Modularity | Efficiency | Transitivity | SWP |
| --- | --- | --- | --- | --- | --- |
| WAB-AQ | 26.62 | 2.11 | -18.51 | -3.81 | -3.55 |
| Factor Sum | 0.84 | -0.05 | -0.66 | 0.07 | -0.05 |
| Lexical | 0.96 | 0.02 | -0.01 | -0.75 | 0.04 |
| Auditory Comp. | -0.45 | -0.33 | 0.01 | 0.13 | -0.10 |
| Phonology | 0.64 | 0.18 | -0.26 | -0.30 | -0.39 |
| Cognitive/Semantic | -0.30 | 0.08 | -0.39 | 0.98 | 0.39 |

Streamline - Whole Brain
| Behavior | Edge Weights | Modularity | Efficiency | Transitivity | SWP |
| --- | --- | --- | --- | --- | --- |
| WAB-AQ | 5.64 | -4.70 | 1.59 | 4.41 | 1.75 |
| Factor Sum | 0.08 | -0.39 | 0.21 | 0.28 | 0.10 |
| Lexical | 0.34 | -0.27 | -0.07 | -0.18 | -0.08 |
| Auditory Comp. | 0.01 | 0.18 | -0.19 | 0.18 | -0.04 |
| Phonology | -0.40 | -0.20 | 0.79 | 0.56 | 0.16 |
| Cognitive/Semantic | 0.14 | -0.11 | -0.31 | -0.28 | 0.06 |

Streamline - Left Hemisphere
| Behavior | Edge Weights | Modularity | Efficiency | Transitivity | SWP |
| --- | --- | --- | --- | --- | --- |
| WAB-AQ | 5.41 | -2.98 | 0.39 | 2.39 | 0.22 |
| Factor Sum | -0.76 | -0.46 | 0.69 | 0.25 | -0.33 |
| Lexical | 0.66 | -0.08 | -0.18 | -0.32 | 0.10 |
| Auditory Comp. | -0.20 | -0.39 | -0.13 | -0.01 | 0.02 |
| Phonology | -0.76 | -0.02 | 0.80 | 0.60 | -0.17 |
| Cognitive/Semantic | -0.45 | 0.03 | 0.19 | -0.02 | -0.27 |

Simulated FA - Whole Brain
| Behavior | Edge Weights | Modularity | Efficiency | Transitivity | SWP |
| --- | --- | --- | --- | --- | --- |
| WAB-AQ | 7.34 | -2.58 | -5.69 | 2.88 | -3.67 |
| Factor Sum | 0.52 | -0.14 | -0.42 | 0.06 | -0.14 |
| Lexical | 0.42 | -0.13 | -0.17 | -0.08 | -0.19 |
| Auditory Comp. | 0.43 | 0.07 | -0.60 | -0.06 | 0.18 |
| Phonology | -0.81 | -0.16 | 0.78 | 0.47 | -0.19 |
| Cognitive/Semantic | 0.48 | 0.07 | -0.43 | -0.27 | 0.06 |

FA - Left Hemisphere
| Behavior | Edge Weights | Modularity | Efficiency | Transitivity | SWP |
| --- | --- | --- | --- | --- | --- |
| WAB-AQ | 15.44 | 5.15 | -9.94 | 2.63 | -1.47 |
| Factor Sum | 0.52 | 0.12 | -0.34 | 0.14 | -0.03 |
| Lexical | 0.45 | 0.18 | -0.11 | 0.03 | 0.14 |
| Auditory Comp. | 0.06 | -0.25 | -0.30 | -0.06 | -0.08 |
| Phonology | 0.28 | 0.06 | -0.07 | -0.04 | -0.29 |
| Cognitive/Semantic | -0.27 | 0.13 | 0.14 | 0.22 | 0.20 |

Streamline - Whole Brain
| Behavior | Edge Weights | Modularity | Efficiency | Transitivity | SWP |
| --- | --- | --- | --- | --- | --- |
| WAB-AQ | 3.85 | -4.99 | 4.10 | 6.30 | 1.72 |
| Factor Sum | 0.00 | -0.41 | 0.33 | 0.37 | 0.11 |
| Lexical | 0.31 | -0.26 | -0.05 | -0.15 | -0.09 |
| Auditory Comp. | -0.15 | 0.14 | 0.02 | 0.31 | -0.04 |
| Phonology | -0.38 | -0.20 | 0.80 | 0.58 | 0.17 |
| Cognitive/Semantic | 0.22 | -0.09 | -0.45 | -0.37 | 0.06 |

Streamline - Left Hemisphere
| Behavior | Edge Weights | Modularity | Efficiency | Transitivity | SWP |
| --- | --- | --- | --- | --- | --- |
| WAB-AQ | 5.07 | -2.85 | 0.72 | 2.72 | 0.10 |
| Factor Sum | -0.79 | -0.46 | 0.72 | 0.27 | -0.33 |
| Lexical | 0.53 | -0.10 | -0.08 | -0.28 | 0.03 |
| Auditory Comp. | -0.18 | -0.39 | -0.14 | 0.00 | 0.04 |
| Phonology | -0.77 | -0.02 | 0.82 | 0.60 | -0.18 |
| Cognitive/Semantic | -0.37 | 0.04 | 0.13 | -0.05 | -0.23 |

*Note:* Beta weights for the simulated models are the means from the simulated permutations. The betas for the WAB-AQ represent the relationship between standardized network measures and raw WAB-AQ values to aid interpretability. The betas for all other measures are between standardized network measures and the factor scores.

